# High-resolution structure of a replication-initiation like configuration of influenza polymerase active site visualises the essential role of a conserved dibasic motif in the PA subunit

**DOI:** 10.1101/2023.11.20.567839

**Authors:** Petra Drncova, Tim Krischuns, Nadia Naffakh, Stephen Cusack

## Abstract

Influenza polymerase, comprising subunits PA, PB1 and PB2, transcribes the negative-sense genomic viral RNA (vRNA) into mRNA or replicates it first into complementary RNA (cRNA) and then back to vRNA. Here we investigate the mechanism of *de novo* (unprimed) initiation of vRNA to cRNA replication. We present a high-resolution structure of A/little-yellow-shouldered-bat/H17N10 polymerase with the 3’ end of the template in the RNA synthesis active site, both in the apo-state and after soaking with GTP and CTP. The priming GTP and incoming CTP are observed to base-pair to template nucleotides C2 and G3 at the -1 and +1 positions respectively, thus representing a replication initiation-like state. This configuration is stabilised by partial stacking of the tip of the priming loop on the GTP:C2 base-pair and the interaction of PB1/H649 and dibasic motif residues PA/R658 and K659 with the triphosphate of the priming GTP. The dibasic motif is universally conserved in orthomyxovirus PA subunits. Trans-complementation assays in cells using mutants of PA/K659 show that the dibasic motif is specifically essential for replication. These results shed light on the mechanism of replication initiation even though vRNA to cRNA replication is expected to be terminally initiated, with priming ATP and incoming GTP base-pairing to template nucleotides U1 and C2 at the -1 and +1 positions respectively, implying a different position of the template.

## Introduction

The heterotrimeric influenza polymerase, comprising subunits PA, PB1 and PB2, both transcribes and replicates the negative sense viral RNA (vRNA) genome in the nucleus of the infected cell (Te Velthuis *et al*, 2021; Wandzik *et al*, 2021). The functional unit of transcription and replication is the vRNP (viral ribonucleoprotein complex) in which the viral RNA is coated by viral nucleoprotein (NP) in a superhelical structure. The polymerase is at one end of the vRNP bound to the so-called promoter, which comprises the conserved, quasi-complementary 3’ and 5’ ends of the vRNA. Whereas transcription generates *bona fide* capped and polyadenylated viral mRNAs, replication generates unmodified full-length copies. The 5’ cap of the viral mRNA originates from initiation of transcription being by a short, capped primer cleaved by the polymerase from nascent pol II transcripts, a process known as cap-snatching (Plotch *et al*, 1981). Polyadenylation occurs by stuttering at a conserved oligo-U sequence proximal to the 5’ end of the vRNA template (Poon *et al*, 1999). The detailed mechanism of transcription by influenza polymerase is now reasonably well-understood thanks to multiple high-resolution structural snapshots at different points along the transcription cycle (Kouba *et al*, 2019; Pflug *et al*, 2018; Reich *et al*, 2014; Wandzik *et al*, 2020). Replication encompasses both complementary RNA (cRNA) synthesis from genomic vRNA and *vice versa*. Crucially, replication is coupled to packaging of the replicate into progeny cRNP or vRNP, this requiring nuclear import of newly synthesised polymerase and NP. Unprimed (‘*de novo*’) vRNA to cRNA synthesis is thought to be terminally initiated by alignment of ATP and GTP with the first two nucleotides at the 3’ end of the vRNA template (1-**UC**GUUU) allowing formation of the dinucleotide pppApG in the first catalytic step (Deng *et al*, 2006). Stabilisation of the replication initiation complex requires the so-called priming loop, a loop element of the PB1 subunit that is adjacent to the active site at the initiation of RNA synthesis (Pflug *et al*, 2014; Te Velthuis *et al*, 2016), necessitating its later extrusion from the active site cavity during the initiation to elongation transition (Kouba *et al*., 2019). In contrast, *de novo* initiation of cRNA to vRNA replication occurs internally at positions 4 and 5 of the cRNA template (1-UCA**UC**U) (Deng *et al*., 2006; Zhang *et al*, 2010) and is not dependent on the priming loop (Te Velthuis *et al*., 2016). The template is then thought to backtrack and the initially formed pppApG realigns to positions 1 and 2 of the template to act as a terminal primer for subsequent vRNA synthesis. Formation of a ‘symmetric’ polymerase dimer (i.e. with a symmetric dimer interface) has been implicated in the realignment step (Chen *et al*, 2019; Fan *et al*, 2019). The current model for co-replicational packaging involves a distinct polymerase dimer with an asymmetric interface between the so-called ‘replicase’, which actively performs the RNA synthesis and an ‘encapsidase’, which nucleates the progeny RNP by binding the emergent 5’ end of the replicate (Carrique *et al*, 2020). The replicase-encapsidase dimer is stabilised by the host protein ANP32, a multifunctional chaperone that is implicated in host restriction of influenza virus replication (Long *et al*, 2016). ANP32 is also proposed to recruit incoming NP to the replication complex to participate in co-replicational packaging by means of a direct interaction between the C-terminal LCAR (low complexity acidic region) of ANP32 and NP (Wang *et al*, 2022). Although significant progress has recently been made in understanding the distinct mechanisms of replication, structures of key steps are missing and hence several outstanding questions remain. These include (a) the exact details of terminal or internal *de novo* initiation for respectively cRNA or vRNA synthesis, including the positioning of the template and the role of the priming loop, (b) the determination of whether or not the translocating template takes the same trajectory into the 3’ end secondary site, as observed for transcription (Wandzik *et al*., 2020), (c) the mechanism by which anti-termination occurs allowing RNA synthesis to proceed through the 5’ hook to the extreme 5’ end of the template, thus circumventing poly-adenylation in the case of vRNA to cRNA replication; and finally, (d) the details of ANP32 mediated NP recruitment and progressive encapsidation of the replicate.

Here we focus on the mechanism of *de novo* replication initiation. We crystallised vRNA promoter-bound A/little-yellow-shouldered-bat/H17N10 (A/bat) polymerase in a form that yields an unusually high-resolution (1.9 Å) structure, which exhibits the 3’ end of the template in the active site adjacent to the priming loop. We obtained a second structure after soaking in two NTPs (priming and initiating) allowing visualisation of a pre-catalytic, replication-like initiation complex. The structure highlighted the role of the priming loop and a conserved basic motif in PA (PA/658-RK in A/bat) in stabilising the triphosphate of the priming NTP. Using a trans-complementation system in a cell-based vRNP reconstitution assay, we showed that the dibasic motif is required for replication, but not transcription. These results are discussed in the light of previous biochemical results on replication initiation that perhaps highlight the difficulty of reproducing genuine replication activity *in vitro* with polymerase alone, although recent progress has been made in this direction (Zhu *et al*, 2023).

## Results

### High resolution crystal form of bat influenza A/H17N10 polymerase

Previously we used a capped primer together with a vRNA promoter with a three nucleotide extension to 3′ end of the template to stabilise and crystallise a transcription-initiation like state of influenza B polymerase (Figure 1a)(Kouba *et al*., 2019). To try to obtain a higher resolution crystal form, a similar strategy was applied to crystallise the transcription initiation state of A/bat polymerase. Crystals of space-group *P*2_1_2_1_2_1_ that diffract slightly anisotropically to a maximum resolution of ∼1.9 Å were obtained (Table 1). This is the highest resolution crystal form so far obtained for a complete influenza polymerase.

**Figure 1.**
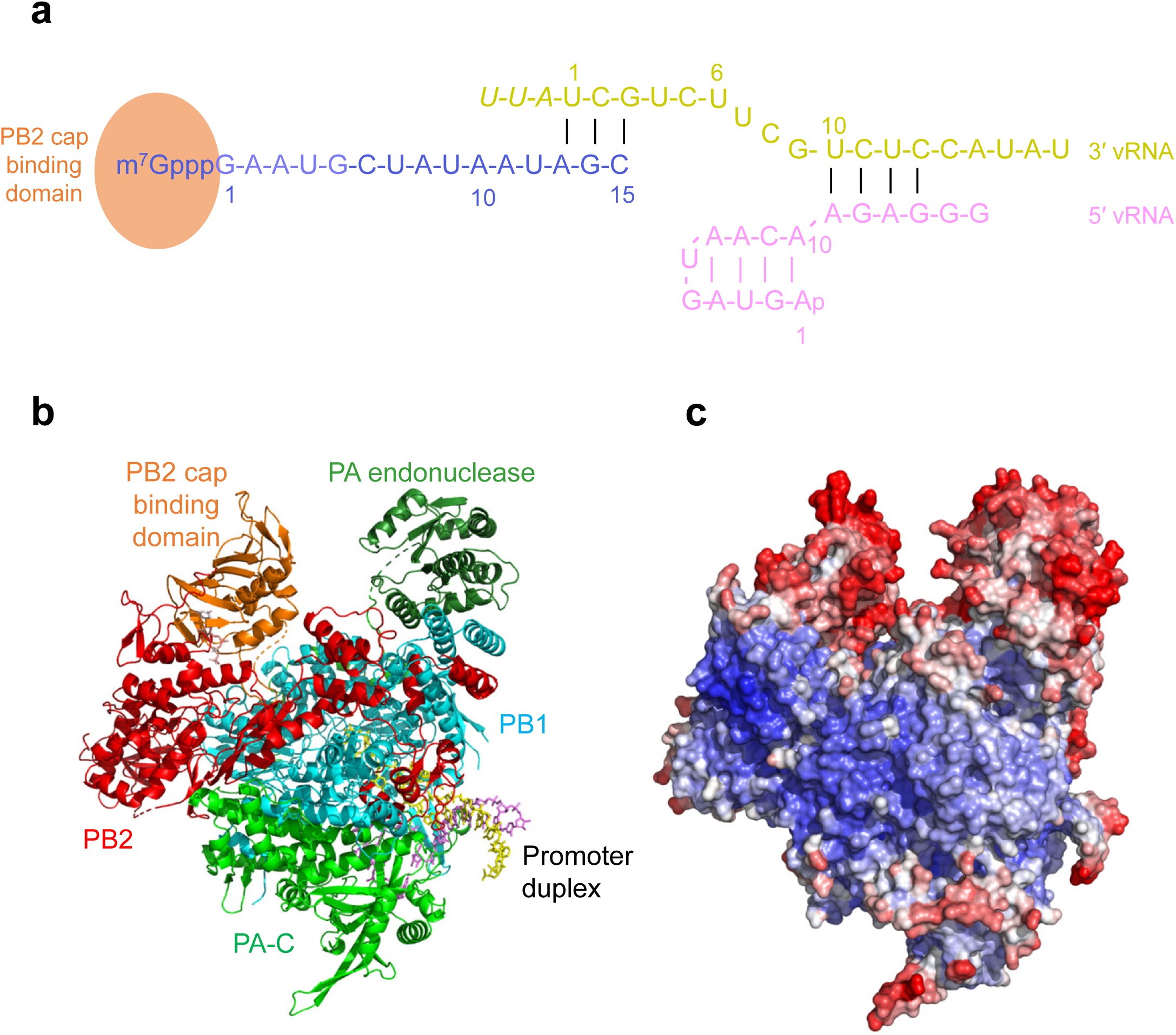
RNAs used in crystallisation and the overall structure. **(a).** Schematic showing the RNAs used to crystallise the putative transcription initiation state of A/bat polymerase. 15-mer capped primer (blue), 21-mer template (yellow, with three extra nucleotides added to the native 3’ end in italics) and 16-mer 5’ vRNA (pink). **(b).** Overall view of the A/bat polymerase crystal structure in ribbon representation, with PA-N (the endonuclease, forest green), PA-C (green), PB1 (cyan), PB2 (red) except cap-binding domain (orange). The promoter 3’ strand is yellow and the 5’ strand is violet. **(c).** Overall view of the A/bat polymerase in surface representation in the same orientation as in (**b**) and coloured according to B-factors (blue-white-red, 20-120 Å^2^). Higher B-factors indicate more disorder.

**Table 1.** Crystallographic data collection and refinement statistics.

|  |  |  |
| --- | --- | --- |
| **Crystal** | Bat A/H17N10 polymerase<br>+3' vRNA 1-18+3<br>+5' vRNA 1-16<br>+14-mer capped RNA primer | Bat A/H17N10 polymerase<br>+3' vRNA 1-18+3<br>+5' vRNA 1-16<br>+14-mer capped RNA primer<br>+GTP+CTP |
| **PDB ID** | 9F2R | 9F37 |
| **Diffraction Data** |  |  |
| Space group | $P2_12_12_1$ | $P2_12_12_1$ |
| Cell dimensions (Å) | a= 91.08 b= 119.27<br>c= 251.56 | a= 90.94 b= 119.01<br>c= 251.38 |
| Wavelength (Å) | 0.9795 | 0.9795 |
| Beamline | Diamond I04 | Diamond I03 |
| No. Crystals | 2 | 1 |
| Resolution range (last shell)<br>(Å) | Ellipsoidal 1.965 Å<br>50.0-1.965 (2.189-1.965) | Ellipsoidal 1.905 Å<br>86.6-1.91 (2.11-1.91) |
| Completeness (last shell) (%) | Overall 69.7<br>Ellipsoidal 95.7 (68.4) | Overall 71.5<br>Ellipsoidal 96.3 (75.6) |
| R-merge (last shell) | 0.159 (1.227) | 0.084 (0.955) |
| R-meas (last shell) | 0.166 (1.290) | 0.091 (1.04) |
| I/σI (last shell) | 9.7 (1.8) | 13.3 (1.8) |
| CC(1/2) (last shell) | 0.997 (0.736) | 0.999 (0.708) |
| Redundancy (last shell) | 13.3 (10.7) | 6.9 (6.4) |
| **Refinement** |  |  |
| Reflections used in<br>refinement: work (free) | 129144 (6693) | 144035 (7590) |
| R-work (last shell) | 0.189 (0.317) | 0.188 (0.331) |
| R-free (last shell) | 0.246 (0.353) | 0.232 (0.409) |
| **Composition** |  |  |
| Total non-hydrogen atoms | 19394 | 19038 |
| Protein residues | 2176 | 2173 |
| RNA | 31 | 32 |
| Ligands | 5: 1x Mg, m<sup>7</sup>GTP, 3x GOL, | 10: GTP, CTP, 2x Mg, m<sup>7</sup>GTP,<br>5x GOL |
| Water | 1093 | 700 |
| **Geometry and B-factors** |  |  |
| RMS(bonds) | 0.005 | 0.006 |
| RMS(angles) | 1.407 | 1.423 |
| Average B-factor (Å<sup>2</sup>) | 47.991 | 49.81 |
| Ramachandran favored (%) | 94.86 | 95.73 |
| Ramachandran outliers (%) | 0.60 | 0.32 |
| Rotamer outliers (%) | 3.29 | 3.87 |
| Molprobity score | 2.02 | 2.07 |
| Clash score | 5.07 | 6.00 |

The asymmetric unit of the crystal contains the polymerase trimer with all domains visible (Figure 1b). However, whereas the core and active site of the polymerase are extremely well-ordered, the peripheral cap-binding and endonuclease domains are poorly ordered, as indicated by the B-factors (Figure 1c). The promoter duplex does not make RNA-RNA crystal contacts, which is unusual for A/bat polymerase crystals (Pflug *et al*., 2018; Pflug *et al*., 2014). Instead the last visible template base (A15) packs against the main-chain of 733-GR-734 of PB2 from a neighbouring polymerase. The high-resolution enables a highly accurate model to be built for much of the polymerase and the vRNA and hundreds of water molecules can be placed (Supp. Figure 1a). The polymerase is in the ‘transcription active’ (transcriptase) configuration (Pflug *et al*., 2018; Thierry *et al*, 2016), but there is no primer-template duplex in the active site and only weak density for the 5′-m^7^Gppp of the primer in the cap-binding site.

### Structure of the RNA synthesis active site with and without initiating NTPs

The high resolution allows unambiguous definition of the path of the template from the duplex region of the promoter into the active site (Supp. Figure 1b). The 3′ end of the template is visible up to the phosphate of U1, but there is no density for the three-nucleotide extension (Supp. Figure 1b). The priming loop is well-ordered and adjacent to the vRNA 3′ end (Figure 2a). As observed in previous structures exhibiting the free vRNA 3’ end in the polymerase active site, template nucleotide C2 is at the -1 position and G3 is at the +1 position (Kouba *et al*., 2019; Pflug *et al*., 2018; Reich *et al*, 2017). Since there is no incoming nucleotide, the motif B loop is in the open state (Kouba *et al*., 2019)

**Figure 2.**
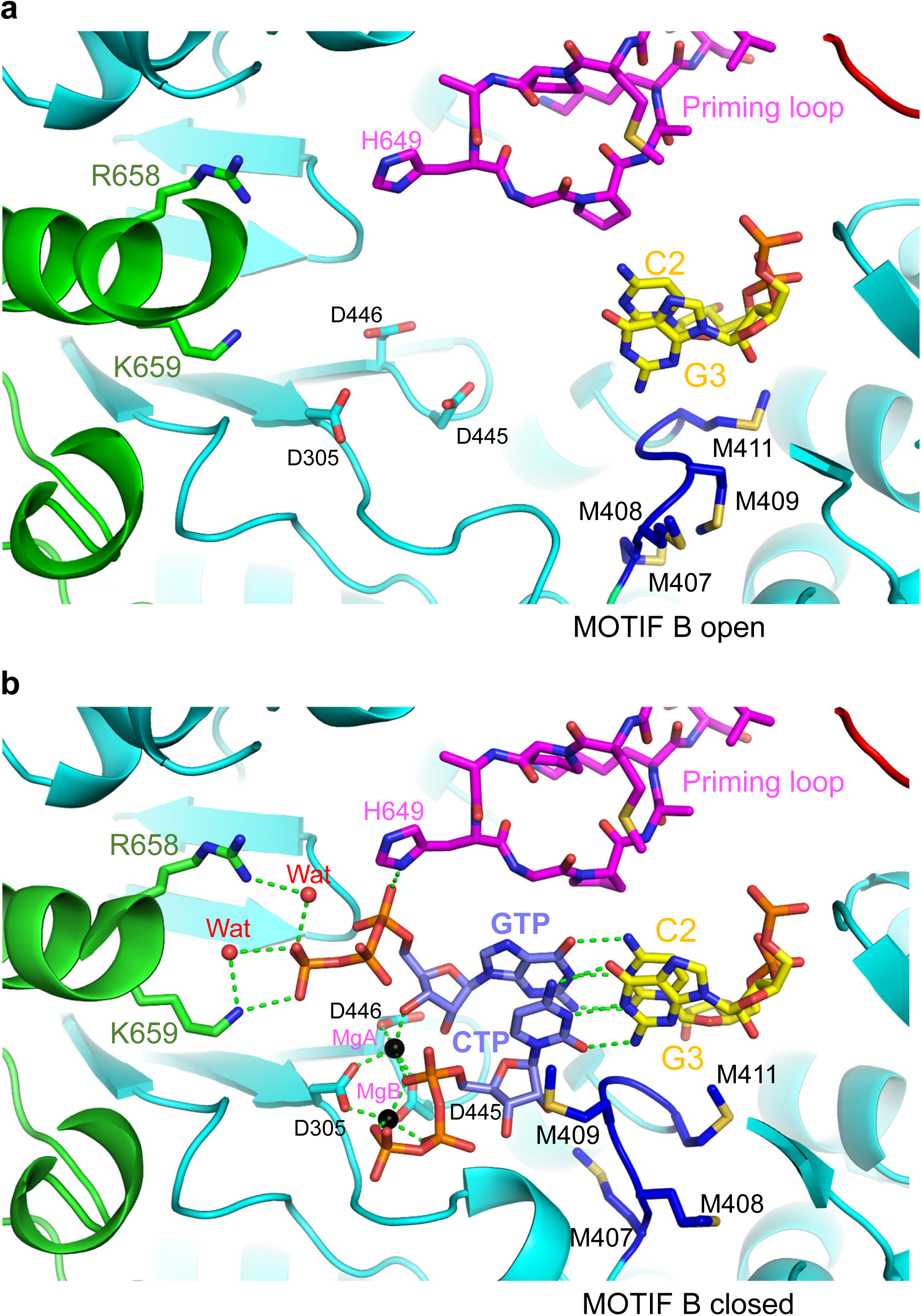
Structural details of the RNA synthesis active site. **(a).** View into the active site of the apo-structure with PA (green), PB1 (cyan) and PB2 (red), 3’ end template nucleotides (yellow), with key residues in stick representation. PB1/D305, D445 and D446 are the active site aspartates. The priming loop (magenta) is fully ordered. There is no incoming NTP and hence the motif B loop PB1/407-MMMGM-411 (blue) is in the open position. **(b).** View into the active site of the GTP+CTP soaked structure in the same orientation and colours as (**a**). The base of U1, which is ordered in this structure, is omitted for clarity. The priming GTP and incoming CTP (slate blue) are at the -1 and +1 positions respectively. The presence of the CTP at the +1 position induces the closed configuration of the motif B loop PB1/407-MMMGM-411 (blue). Two fully co-ordinated magnesium ions are bound at the catalytic center. The PA dibasic motif PA/R658 and K659, together with H649 from the priming loop and two water molecules (red spheres) stabilise the triphosphate of the priming GTP. Priming loop residues 650G and 651P partially stack on the GTP:C2 base-pair at the -1 position. Hydrogen bonds and metal co-ordination are in green dotted lines.

The presence of the free template 3′ end in the active site suggested that incoming, initiating NTPs could be visualized by soaking. The crystallization conditions are promising for this, as they do not contain competing phosphate, unlike previous polymerase crystal forms. Since in the crystal structure, template bases C2 and G3 are at respectively the -1 and +1 positions we soaked GTP and CTP to mimic a *de novo* replication-initiation state. This resulted in a second high-resolution dataset (Table 1). The highly ordered active site now shows GTP and CTP forming base-pairs with the template in respectively the -1 and +1 positions (Supp. Figure 1a, Figure 2b) and additionally the base and ribose of U1 become visible, but still not the three nucleotide extension (Supp. Figure 1b). This is the first structure showing a priming NTP in the -1 position. It reveals the role of the two basic residues PA/658-RK and PB1/H649 from the priming loop, together with water molecules, in stabilising the position of the triphosphate of the priming GTP (Figure 2b). In addition, priming loop residues PB1/650G and 651P partially stack on the GTP:C2 base-pair at the -1 position. There are two well-defined magnesium ions co-ordinated by the three active site aspartates (PB1/D305, D445 and D446) and the triphosphate of the CTP in an apparently catalytic configuration (Figure 2b). However, perhaps due to crystal constraints, the reaction does not occur i.e. the substrate complex with GTP and CTP is observed, not the product complex, which would have pppGpC + PPi (pyrophosphate). Since there is CTP in the +1 incoming nucleotide position, the motif B loop undergoes an induced fit flip to the closed conformation with PB1/M409 stacking under the CTP (Figure 2b), consistent with what has been previously observed for influenza B polymerase (Kouba *et al*., 2019).

### The conserved dibasic motif in PA is specifically required for replication

The structure with incoming NTPs shows that there are direct and water-mediated contacts between PA/658-RK and PB1/H649 and the triphosphate of the priming GTP. The dibasic motif PA/658-RK in A/bat (PA/663-RK in human/avian FluA, PA/659-RR in FluB) is very highly conserved in the PA subunit of orthomyxovirus polymerases, with very exceptionally a methionine at the second position in some Quaranjaviruses (Figure 3a). This motif, also occurring in bunyavirus L proteins, has previously been designated as polymerase motif G (Gerlach *et al*, 2015).

**Figure 3.**
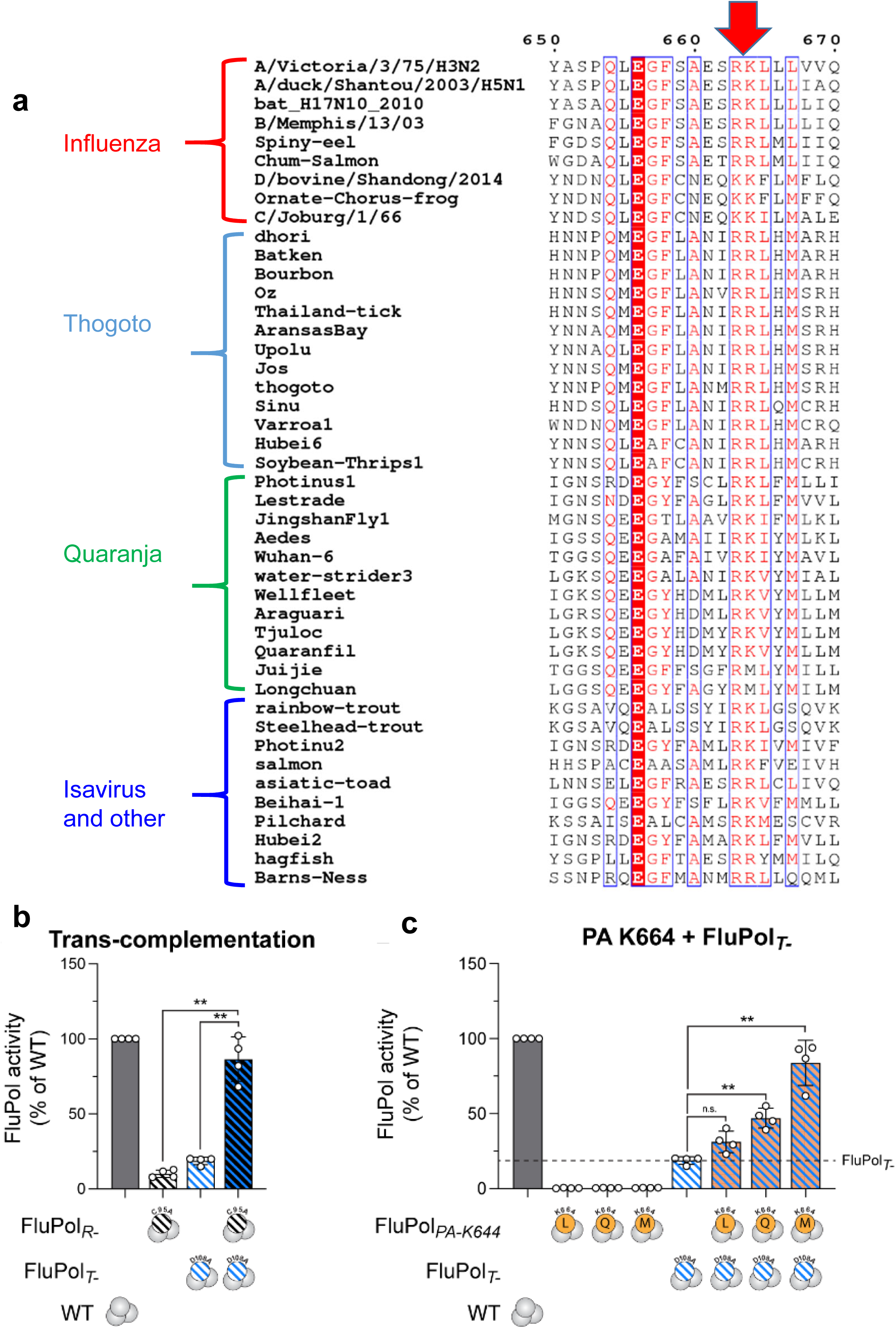
The PA dibasic motif and cell-based trans-complementation assay. **(a)** Alignment of PA sequences, from diverse orthomyxoviruses highlighting the conservation of the dibasic motif (red arrow). Residue numbers refer to the top sequence. The alignment was done with ClustalW (Larkin *et al*, 2007) and visualized with ESPRIPT (Gouet *et al*, 1999). **(b)** Trans-complementation of a replication-defective (PA/C95A, black, (Nilsson-Payant *et al*., 2018)) and a transcription-defective (PA/D108A, blue, (Hara *et al*., 2006)) FluPol was measured by vRNP reconstitution in HEK-293T cells upon transient transfection, using a model vRNA encoding the Firefly luciferase. Firefly luminescence was measured at 48 hours post-transfection and normalised to a Renilla luciferase transfection control. The data are represented as percentage of PA wild-type (dark grey). **p < 0.002 (one-way repeated measure ANOVA; Dunnett’s multiple comparisons test). **(c)** The FluPol activity of the indicated PA/K664 mutants (orange) were assessed either alone or in a FluPol trans-complementation assay by co-expression of a transcription-deficient FluPol (PA/D108A, (Hara *et al*., 2006)) as in (A). **p < 0.002 (one-way repeated measure ANOVA; Dunnett’s multiple comparisons test). n.s. : non-significant. The background signal is represented by a dotted line.

To test the importance of the PA dibasic motif in replication initiation, we used a vRNP reconstitution assay with strain A/WSN/33 (H1N1). Here, polymerase activity is measured upon transient co-transfection of plasmids encoding all vRNP components including a viral-like RNA encoding a firefly luciferase. In this assay, reporter activity depends on FluPol replication as well as transcription activity. In particular, we made use of the observation that FluPol transcription and replication can be decoupled by using either a replication-deficient PA/C95A mutant (black) (Nilsson-Payant *et al*, 2018) or a transcription-deficient PA/D108A mutant (blue) (Hara *et al*, 2006). PA/C95A and PA/D108A individually only had residual activity but almost reached wild-type levels upon co-transfection (Figure 3b) as previously described (Nilsson-Payant *et al*., 2018).

The first basic residue is in most cases an arginine (Figure 3a), and the structure shows that it has a significant structural role in stabilising the local protein environment in addition to co-coordinating a water molecule to the priming triphosphate. Therefore, this residue was not mutated as it could simply abolish activity due to a too great structural disturbance of the active site. The second residue (PA/K664 in A/H1N1, PA/K659 in A/bat), which exceptionally in some viruses is a methionine (Figure 3a), was mutated to a hydrophobic residue (L, M) or a glutamine (Q). All three investigated PA K664 mutants individually abolished polymerase activity as examined in a standard vRNP reconstitution assay (Figure 3c). To test whether these mutants have a specific replication defect, trans-complementation by the transcription-defective/replication-competent mutant PA/D108, was assessed. Whereas PA/K664L and PA/K664Q could be partially complemented by PA/D108A, PA/K664M levels nearly reached wild-type levels upon-transfection (Figure 3c). This shows that the investigated PA/K66M mutants are specifically replication deficient but transcription competent. Taken together, the high-resolution structure and the complementation assays strongly suggest that the PA dibasic motif is essential for replication initiation due to its role in stabilising the configuration of the priming NTP.

## Discussion

We describe a new high-resolution crystal form of the A/bat polymerase, in which the 3’ end of the template is bound in the RNA synthesis active site. As also observed for FluB polymerase by X-ray crystallography (Reich *et al*., 2017) and for A/H7N9 polymerase by cryo-electron microscopy (Krischuns *et al*, 2024), the 3’ end is aligned with C2 at the -1 position and G3 at the +1 position, adjacent to the tip of the priming loop, with U1 partially disordered (Figure 4a). This allows binding of initiating nucleotides GTP and CTP, stabilised by essential interactions with the PA dibasic motif and the priming loop, which can then react to give pppGpC through the catalytic action of the polymerase. RNA synthesis, mimicking vRNA to cRNA replication, would thus proceed by translocation and addition of the next nucleotide ATP. Unprimed initiation of the vast majority of products with pppGpC has been confirmed by next generation sequencing in the case of A/H7N9 polymerase (Krischuns *et al*., 2024). Even though this seems to be the ‘natural’ unprimed RNA synthesis activity of influenza polymerase, at least *in vitro*, it does not correspond with the generally accepted view that vRNA to cRNA replication is terminally initiated with production or pppApG from ATP and GTP (Figure 4b), thus ensuring a full-length genome copy. However, this terminal initiation would require the 3’ end to backtrack by one position, aligning U1 at the -1 position and C2 at the +1 position. There would then be seven nucleotides connecting the +1 position to the promoter duplex, instead of six, which could be accommodated structurally, but has never been observed so far. Consistent with this, soaking of ATP and GTP (or ApG) into the crystals did not reveal any bound nucleotide, neither does GTP alone bind strongly, whereas CTP is observed to bind alone at the +1 position (data not shown), with in none of these cases the template changing alignment.

**Figure 4.**
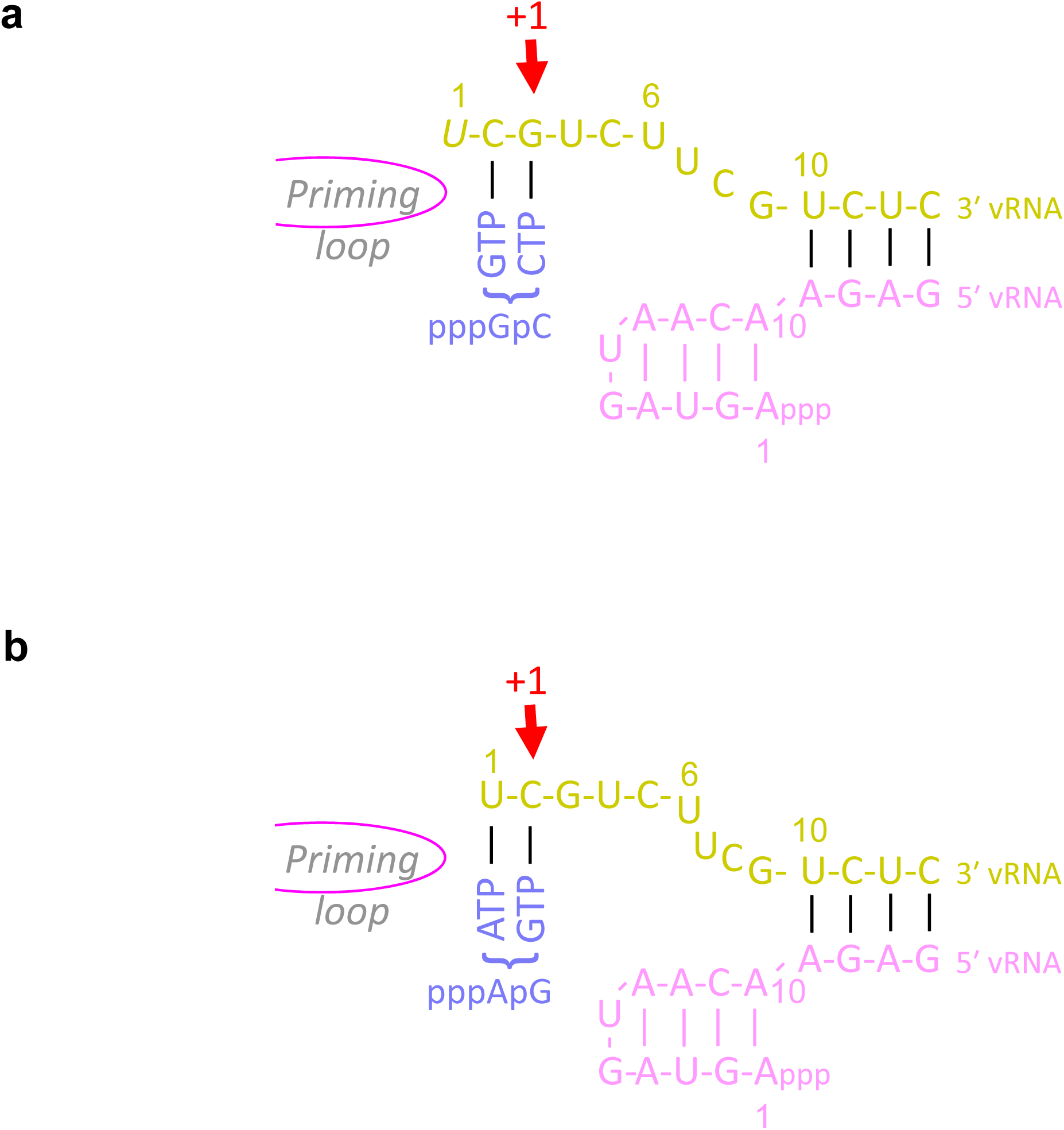
Schematics diagrams showing *de novo* vRNA to cRNA replication. **(a)** Template alignment observed in the crystal structure corresponding to internal *de novo* initiation with GTP and CTP base-pairing with C2 and G3 of the template at respectively the -1 and +1 positions. There are six nucleotides connecting the +1 position to the promoter duplex. **(b)** Template alignment corresponding to terminal *de novo* initiation with ATP and GTP base-pairing with U1 and C2 of the template at respectively the -1 and +1 positions. There are seven nucleotides connecting the +1 position to the promoter duplex.

Previous biochemical studies are ambiguous about this issue. Internal initiation at position 2 of vRNA has been previously described (Vreede & Brownlee, 2007; Zhang *et al*., 2010). In the (Vreede & Brownlee, 2007) study, it was found that pre-expressed viral polymerase and NP binds full-length cRNA (i.e. with 5’ terminal A) synthesized by infecting virions in the absence of transcription (i.e. cycloheximide present), whereas vRNPs purified from virions mostly produced cRNAs lacking the first A. However, the later could perhaps be ascribed to the fact that many vRNAs from purified virus were found to lack a 3’ terminal U (Vreede & Brownlee, 2007). In the (Zhang *et al*., 2010) study using highly purified recombinant polymerase, pppGpC is systematically observed, which corresponds most closely to what we find, and the authors propose that the terminal A might be subsequently added by a cellular poly(A) polymerase. On the other hand, a third study, using partially purified polymerase, report pppApG formation at position 1 for vRNA and internally at position 4 for cRNA (Deng *et al*., 2006). Thus although it is logical to assume that to generate a full-length cRNA product, vRNA to cRNA replication is initiated by pppApG formation, exactly how this achieved structurally and mechanistically is still unclear. The suggestion from the experiments reported so far, including the current work, is that it is difficult, and maybe impossible, to mimic terminal vRNA to cRNA replication *in vitro* using pure polymerase alone. Rather, transcription-like activity is default in this situation. Indeed, the observed position of the template is optimal for primed RNA synthesis with either capped RNA ending in -ApG(pC) or simply the dinucleotide ApG. Other factors may be required to induce the template to take a different path into the active site that places U1 at the -1 position. These could include the RNP context and/or viral or host components known to be relevant for replication and product encapsidation, such as newly synthesized NP and polymerase, the essential host factor ANP32 (Carrique *et al*., 2020; Long *et al*., 2016) or the CTD of Pol II (Krischuns *et al*., 2024). Alternatively, phosphorylation, for instance of PB1/S673, could be a mechanism to suppress transcription in favour of replication, perhaps by remodeling the template entrance channel (Dawson *et al*, 2020).

In conclusion, although the exact active site and template configuration that allows terminal initiation of vRNA to cRNA replication remains elusive, this work reveals the key role played by the highly conserved PA dibasic motif in stabilising the priming nucleotide during replication initiation.

## Methods

### Protein purification

A/little-yellow-shouldered-bat/H17N10 polymerase was expressed in insect cells and purified as previously described (Pflug *et al*., 2014; Wandzik *et al*., 2020).

### Crystallisation

Long prism shaped crystals (dimensions ∼300 x 30 µM) were obtained with conditions: 5 mg/ml polymerase with 5.1 mg/ml (19.5 μM) with 1.14 x molar excess of each RNA (v5’ 1-16, v3’ 1-18+3 and 15-mer capped primer, see Figure 1a) mixed in 1:1 ratio of 100 mM amino acids, 100 mM Tris/Bicine pH8.5, 8% ethylene glycol (v/v), 4% PEG 8000 (w/v) by hanging drop at room temperature. Soaking was performed with 5 mM GTP, 5 mM CTP and 5 mM MgCl_2_ for 5h. Crystals were cryo-protected with either an additional 12% ethylene glycol or 25% glycerol before X-ray data collection. The native crystals are of space-group *P*2_1_2_1_2_1_ and diffract slightly anisotropically to a maximum resolution of ∼1.90 Å (Table 1). This is the highest resolution crystal form so far obtained for a complete influenza polymerase.

### RNA sequences

15-mer capped primer (Trilink): m^7^GpppGAAUGCUAUAAUAGC

16-mer 5’ vRNA (IBA): 5’-pAGUAGUAACAAGAGGG

18+3-mer 3’ vRNA (IBA) : 5’-UAUACCUCUGCUUCUGCUAUU

### Structure determination and refinement

Diffraction data were collected on Diamond synchrotron beamlines I03 and I04. Anisotropic data were integrated with AUTOPROC/STARANISO using an ellipsoidal mask with resolution cut-off criteria local (I/sigI) = 1.2 (Tickle *et al*, 2018) (Table 1). Crystal structures were solved by molecular replacement with PHASER (McCoy *et al*, 2007), using a previous A/bat polymerase structure (PDB: 6EVJ) (Pflug *et al*., 2018), rebuilt with COOT (Emsley & Cowtan, 2004) and refined using REFMAC5 (Murshudov, 1997) and Phenix (Afonine *et al*, 2012). Validation was performed using Molprobity (Chen *et al*, 2010) as implemented in the Phenix. Structure figures were produced with Pymol (Schrodinger & DeLano, 2002).

### Plasmids

The A/WSN/33 pcDNA3.1-PB2, -PB1, -PA and pCI-NP plasmids were described previously (Krischuns *et al*, 2022; Lukarska *et al*, 2017). For vRNP reconstitution assays, a pPolI-Firefly plasmid encoding the Firefly luciferase sequence in negative polarity flanked by the 5’ and 3’ non-coding regions of the IAV NS segment was used (Krischuns *et al*., 2022) and the pTK-Renilla plasmid (Promega) was used as an internal control. All mutations were introduced by an adaptation of the QuikChange site-directed mutagenesis (Agilent Technologies) protocol (Zheng *et al*, 2004). Primer and plasmid sequences are available upon request.

### FluPol trans-complementation assay

HEK-293T cells were seeded in 96-well white plates (Greiner Bio-One) the day before transfection. Cells were co-transfected with plasmids encoding the vRNP protein components (PB2, PB1, PA and NP), a pPolI-Firefly plasmid encoding a negative-sense viral-like RNA expressing the Firefly luciferase and the pTK-Renilla plasmid (Promega) as an internal control. Equal amounts of expression plasmids expressing the trans-complementing PA mutants were co-transfected. Firefly and Renilla luciferase activities were measured in duplicates at 48 hpt using the Dual-Glo Luciferase Assay system (Promega) according to the manufacturer’s instructions.

## Acknowledgments

We thank the staff of the Diamond synchrotron for remote access to X-ray beamlines and the EMBL eukaryotic expression (EEF) and high-throughput crystallisation (HTX) facilities within the EMBL-ESRF-ILL-IBS Partnership for Structural Biology. This work was partly supported by ANR grant (ANR-18-CE11-0028) to SC and NN. TK was funded by the ANR grants ANR-18-CE18-0028 and ANR-10-LABX-62-IBEID.

## Author contributions

P.D. expressed, purified and crystallized bat/H17N10 polymerase. S.C. conceived and supervised the project, collected crystallographic data, did the structural analysis and wrote the manuscript with input from all other authors. TK performed in complementation assays, supervised by NN.

## Data Availability

X-ray structures and structure factors are available at PDB: 9F2R (promoter bound) and PDB: 9F37 (promoter bound with GTP and CTP).

## Request for Materials and Correspondence

should be addressed to Stephen Cusack.

---

**Supplementary Figure 1.**
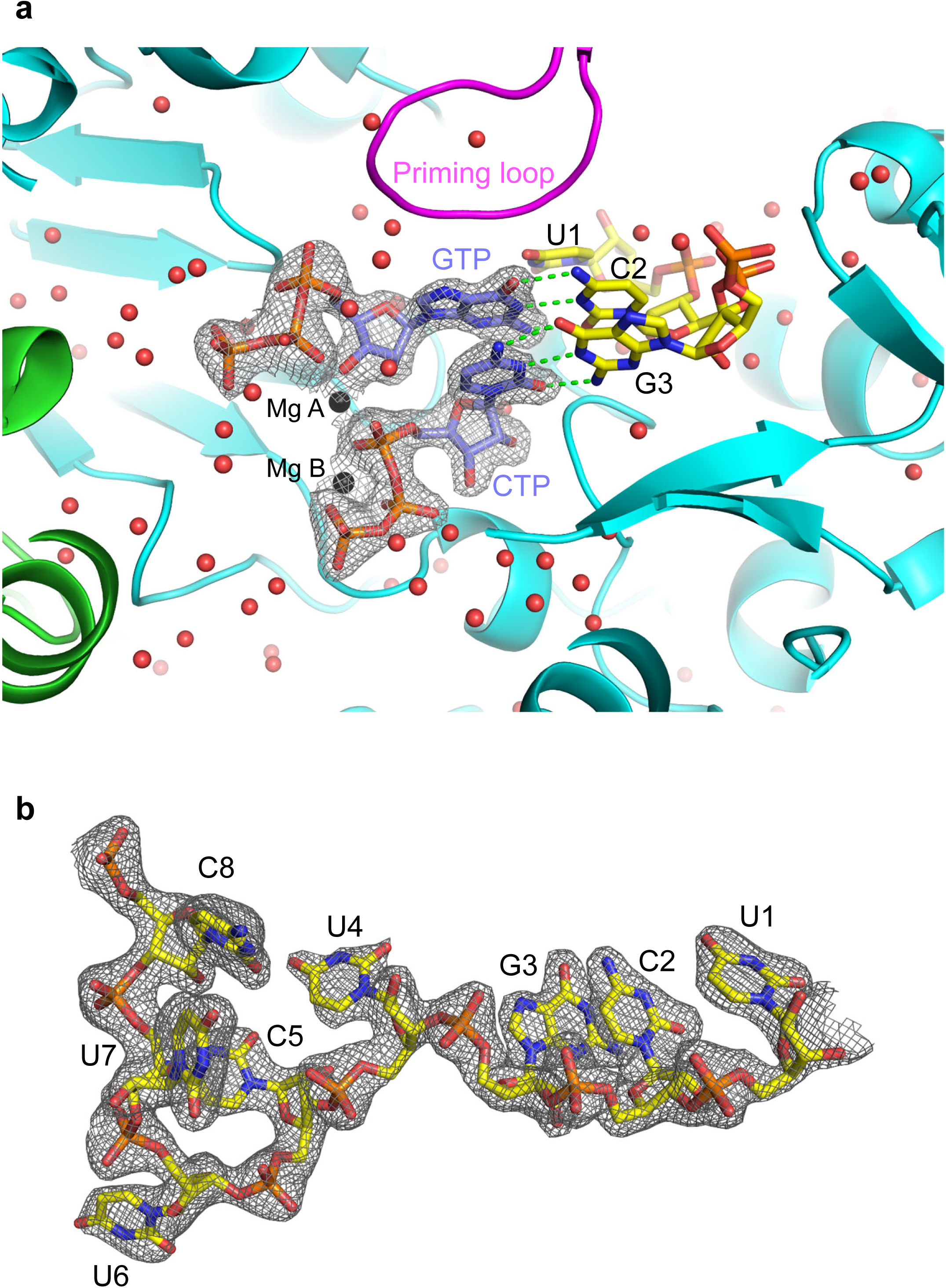
Electron density within the GTP and CTP soaked crystals. **(a)** Unbiased Fo-Fc difference electron density for soaked GTP and CTP (slate blue) within the active site of A/bat polymerase with PA (green), PB1 (cyan), priming loop (magenta) and 3’ end template nucleotides (yellow). The red spheres are water molecules. **(b)** Final 2Fo-Fc electron density for the 3’ end of the template.

## References

Afonine PV, Grosse-Kunstleve RW, Echols N, Headd JJ, Moriarty NW, Mustyakimov M, Terwilliger TC, Urzhumtsev A, Zwart PH, Adams PD (2012) Towards automated crystallographic structure refinement with phenix.refine. *Acta crystallographica Section D*, Biological crystallography 68: 352–367

Carrique L, Fan H, Walker AP, Keown JR, Sharps J, Staller E, Barclay WS, Fodor E, Grimes JM (2020) Host ANP32A mediates the assembly of the influenza virus replicase. Nature 587: 638–643

Chen KY, Santos Afonso ED, Enouf V, Isel C, Naffakh N (2019) Influenza virus polymerase subunits co-evolve to ensure proper levels of dimerization of the heterotrimer. PLoS pathogens 15: e1008034

Chen VB, Arendall WB, 3rd, Headd JJ, Keedy DA, Immormino RM, Kapral GJ, Murray LW, Richardson JS, Richardson DC (2010) MolProbity: all-atom structure validation for macromolecular crystallography. Acta crystallographica Section D, Biological crystallography 66: 12–21

Dawson AR, Wilson GM, Freiberger EC, Mondal A, Coon JJ, Mehle A (2020) Phosphorylation controls RNA binding and transcription by the influenza virus polymerase. PLoS pathogens 16: e1008841

Deng T, Vreede FT, Brownlee GG (2006) Different de novo initiation strategies are used by influenza virus RNA polymerase on its cRNA and viral RNA promoters during viral RNA replication. Journal of virology 80: 2337–2348

Emsley P, Cowtan K (2004) Coot: model-building tools for molecular graphics. *Acta crystallographica Section D*, Biological crystallography 60: 2126–2132

Fan H, Walker AP, Carrique L, Keown JR, Serna Martin I, Karia D, Sharps J, Hengrung N, Pardon E, Steyaert J et al (2019) Structures of influenza A virus RNA polymerase offer insight into viral genome replication. Nature 573: 287–290

Gerlach P, Malet H, Cusack S, Reguera J (2015) Structural Insights into Bunyavirus Replication and Its Regulation by the vRNA Promoter. Cell 161: 1267–1279

Gouet P, Courcelle E, Stuart DI, Metoz F (1999) ESPript: analysis of multiple sequence alignments in PostScript. Bioinformatics 15: 305–308

Hara K, Schmidt FI, Crow M, Brownlee GG (2006) Amino acid residues in the N-terminal region of the PA subunit of influenza A virus RNA polymerase play a critical role in protein stability, endonuclease activity, cap binding, and virion RNA promoter binding. Journal of virology 80: 7789–7798

Kouba T, Drncova P, Cusack S (2019) Structural snapshots of actively transcribing influenza polymerase. Nature structural & molecular biology 26: 460–470

Krischuns T, Arragain B, Isel C, Paisant S, Budt M, Wolff T, Cusack S, Naffakh N (2024) The host RNA polymerase II C-terminal domain is the anchor for replication of the influenza virus genome. Nat Commun 15: 1064

Krischuns T, Isel C, Drncova P, Lukarska M, Pflug A, Paisant S, Navratil V, Cusack S, Naffakh N (2022) Type B and type A influenza polymerases have evolved distinct binding interfaces to recruit the RNA polymerase II CTD. PLoS pathogens 18: e1010328

Larkin MA, Blackshields G, Brown NP, Chenna R, McGettigan PA, McWilliam H, Valentin F, Wallace IM, Wilm A, Lopez R et al (2007) Clustal W and Clustal X version 2.0. Bioinformatics 23: 2947–2948

Long JS, Giotis ES, Moncorge O, Frise R, Mistry B, James J, Morisson M, Iqbal M, Vignal A, Skinner MA et al (2016) Species difference in ANP32A underlies influenza A virus polymerase host restriction. Nature 529: 101–104

Lukarska M, Fournier G, Pflug A, Resa-Infante P, Reich S, Naffakh N, Cusack S (2017) Structural basis of an essential interaction between influenza polymerase and Pol II CTD. Nature 541: 117–121

McCoy AJ, Grosse-Kunstleve RW, Adams PD, Winn MD, Storoni LC, Read RJ (2007) Phaser crystallographic software. J Appl Crystallogr 40: 658–674

Murshudov GN (1997) Refinement of macromolecular structures by the maximum-likelihood method. *Acta crystallographica Section D*, Biological crystallography 53: 240–255

Nilsson-Payant BE, Sharps J, Hengrung N, Fodor E (2018) The Surface-Exposed PA(51-72)-Loop of the Influenza A Virus Polymerase Is Required for Viral Genome Replication. Journal of virology 92

Pflug A, Gaudon S, Resa-Infante P, Lethier M, Reich S, Schulze WM, Cusack S (2018) Capped RNA primer binding to influenza polymerase and implications for the mechanism of cap-binding inhibitors. Nucleic acids research 46: 956–971

Pflug A, Guilligay D, Reich S, Cusack S (2014) Structure of influenza A polymerase bound to the viral RNA promoter. Nature 516: 355–360

Plotch SJ, Bouloy M, Ulmanen I, Krug RM (1981) A unique cap(m7GpppXm)-dependent influenza virion endonuclease cleaves capped RNAs to generate the primers that initiate viral RNA transcription. Cell 23: 847–858

Poon LL, Pritlove DC, Fodor E, Brownlee GG (1999) Direct evidence that the poly(A) tail of influenza A virus mRNA is synthesized by reiterative copying of a U track in the virion RNA template. Journal of virology 73: 3473–3476

Reich S, Guilligay D, Cusack S (2017) An in vitro fluorescence based study of initiation of RNA synthesis by influenza B polymerase. Nucleic acids research 45: 3353–3368

Reich S, Guilligay D, Pflug A, Malet H, Berger I, Crepin T, Hart D, Lunardi T, Nanao M, Ruigrok RW et al (2014) Structural insight into cap-snatching and RNA synthesis by influenza polymerase. Nature 516: 361–366

Schrodinger L, DeLano W (2002) PyMOL Molecular Graphics System. *Available online at* http://wwwpymolorg/pymol

Te Velthuis AJ, Robb NC, Kapanidis AN, Fodor E (2016) The role of the priming loop in influenza A virus RNA synthesis. Nature microbiology 1: 16029

Te Velthuis AJW, Grimes JM, Fodor E (2021) Structural insights into RNA polymerases of negative-sense RNA viruses. Nature reviews Microbiology 19: 303–318

Thierry E, Guilligay D, Kosinski J, Bock T, Gaudon S, Round A, Pflug A, Hengrung N, El Omari K, Baudin F et al (2016) Influenza Polymerase Can Adopt an Alternative Configuration Involving a Radical Repacking of PB2 Domains. Molecular cell 61: 125–137

Tickle IJ, Flensburg C, Keller P, Paciorek W, Sharff A, Vonrhein C, Bricogne G (2018) STARANISO. Global Phasing Ltd Cambridge, United Kingdom

Vreede FT, Brownlee GG (2007) Influenza virion-derived viral ribonucleoproteins synthesize both mRNA and cRNA in vitro. Journal of virology 81: 2196–2204

Wandzik JM, Kouba T, Cusack S (2021) Structure and Function of Influenza Polymerase. Cold Spring Harb Perspect Med 11

Wandzik JM, Kouba T, Drncova P, Karuppasamy M, Pflug A, Provaznik J, Azevedo N, Cusack S (2020) A structure-based model for the complete transcription cycle of influenza polymerase. Cell

Wang F, Sheppard CM, Mistry B, Staller E, Barclay WS, Grimes JM, Fodor E, Fan H (2022) The C-terminal LCAR of host ANP32 proteins interacts with the influenza A virus nucleoprotein to promote the replication of the viral RNA genome. Nucleic acids research 50: 5713–5725

Zhang S, Wang J, Wang Q, Toyoda T (2010) Internal initiation of influenza virus replication of viral RNA and complementary RNA in vitro. The Journal of biological chemistry 285: 41194–41201

Zheng L, Baumann U, Reymond JL (2004) An efficient one-step site-directed and site-saturation mutagenesis protocol. Nucleic acids research 32: e115

Zhu Z, Fan H, Fodor E (2023) Defining the minimal components of the influenza A virus replication machinery via an in vitro reconstitution system. PLoS Biol 21: e3002370

